## Supplementary figures for "Stimulus-specific plasticity in human visual gamma-band activity and functional connectivity"

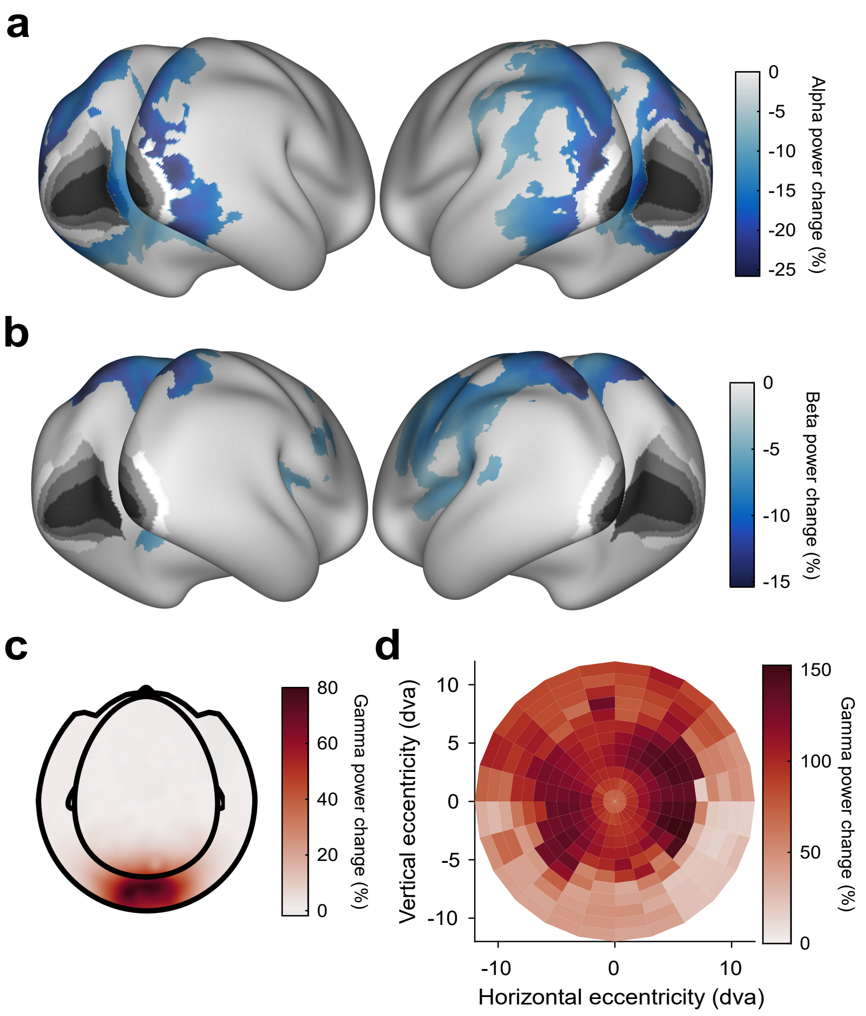


***Figure S1*. Stimulus-induced power changes. Related to Figure 2.** (A) Average stimulus-induced alpha-power change (at individual alpha frequency), source-localized to all cortical dipoles. Values are significance-masked using a $t_{max}$-corrected paired permutation test. Black-to-white shading indicates areas V1, V2, V3, V3A, and V4, as in Fig. 2. Note that alpha-power changes are absent in those areas. (B) Same as (A), but for power changes at the individual beta frequency. (C) Average stimulus-induced gamma-power change on the sensor level. (D) Gamma-power changes (color code) as a function of cortical retinotopic representation (x- and y-axis). Gamma-power changes were first projected into source dipoles. For each dipole in area V1, a group-average retinotopic atlas aligned to the HCP-MMP 1.0 template brain[67] was used to find the corresponding retinotopic position. Subsequently, gamma-power changes were projected from source space into retinotopic space. Gamma-power changes were strongest at the representation of the parafoveal eccentricities.

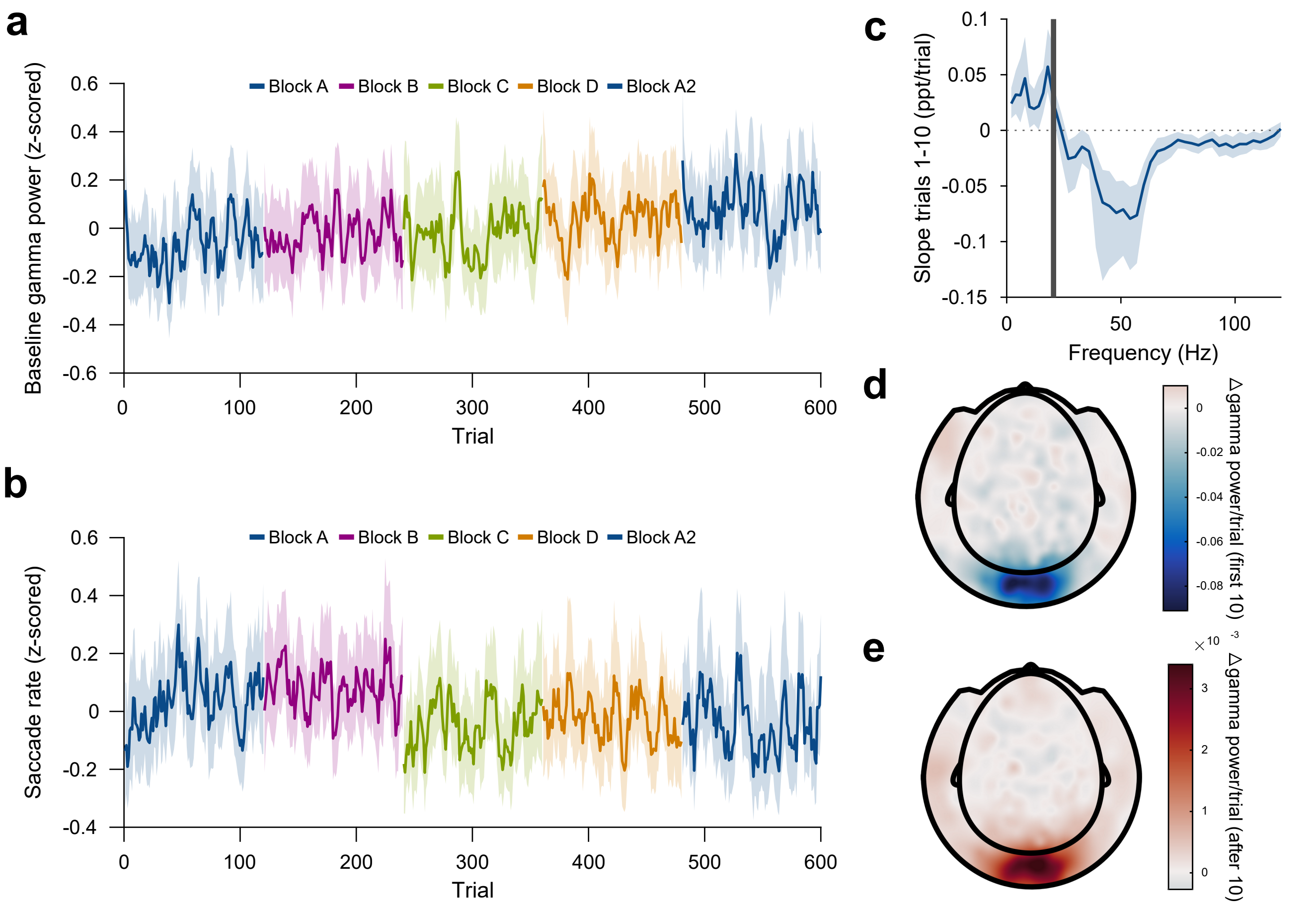


***Figure S2*. Control analyses of repetition-induced gamma changes. Related to Figure 3.** (A) Same as Fig. 3A, but for gamma power in the baseline period (last second before stimulus onset). Baseline gamma power showed an increase with overall trial number, but there was no effect of stimulus-specific repetition number. (B) Same as Fig. 3A, but for microsaccade rate. Microsaccade rate showed a decrease with overall trial number, but there was no effect of stimulus-specific repetition number. (C) Same as Fig. 3E, but for the early trials (trials 1-10). For each frequency, a linear regression across repetitions was fit to the per-trial visually-induced power change in V1/V2 during the early trials (trials 1-10). Average slope and 95% bootstrap CI over subjects is shown. (D) Spatial distribution of the early repetition-related gamma-power decrease: For each MEG gradiometer, a regression line was fit over stimulus repetitions 1-10. Subject-averaged slopes are shown. (E) Same as (D), but for repetitions 11-120.

**a**

**Gamma power model**

**Fixed effects**

| *Variable* | *Estimate* | *SE* | *p* | *2.5%* | *97.5%* |
|---|---|---|---|---|---|
| Intercept | 142.00 | 20.58 | 3.70E-08 | 101.24 | 183.16 |
| Repetition number | 0.40 | 0.03 | 2.00E-16 | 0.33 | 0.46 |
| Trial number | 0.07 | 0.01 | 2.00E-16 | 0.06 | 0.09 |
| Repetition block | 7.80 | 3.45 | 0.024 | 0.91 | 14.87 |
| ITI (s) | 8.13 | 1.63 | 6.27E-07 | 4.96 | 11.35 |
| Microsaccade rate | 0.81 | 0.59 | 0.170 | -0.36 | 1.94 |
| Pupil constriction | 5.67 | 1.97 | 4.05E-03 | 1.82 | 9.45 |
| Early repetitions | 30.79 | 3.99 | 1.34E-14 | 22.37 | 38.75 |

**Random effects**

| *Variable* | *SD* | *2.5%* | *97.5%* |
|---|---|---|---|
| Subject | 98.86 | 74.84 | 123.80 |
| Orientation | 15.12 | 3.49 | 26.06 |
| Residual | 110.34 | 108.99 | 111.58 |

**b**

**Gamma frequency model**

**Fixed effects**

| *Variable* | *Estimate* | *SE* | *p* | *2.5%* | *97.5%* |
|---|---|---|---|---|---|
| Intercept | 34.82 | 2.00 | 2.00E-16 | 30.63 | 38.58 |
| Repetition number | 0.05 | 5.46E-03 | 2.00E-16 | 0.04 | 0.06 |
| Trial number | -0.01 | 1.34E-03 | 3.20E-15 | -0.01 | -0.01 |
| Repetition block | 3.21 | 0.57 | 1.91E-08 | 2.10 | 4.33 |
| ITI (s) | 1.61 | 0.10 | 1.82E-09 | 1.08 | 2.17 |
| Microsaccade rate | 0.03 | 0.33 | 0.778 | -0.16 | 0.23 |
| Pupil constriction | 3.04 | 0.27 | 2.00E-16 | 2.38 | 3.65 |
| Early repetitions | 0.88 | 0.67 | 0.19 | -0.39 | 2.14 |

**Random effects**

| *Variable* | *SD* | *2.5%* | *97.5%* |
|---|---|---|---|
| Subject | 8.70 | 6.43 | 10.91 |
| Orientation | 1.23 | 0.00 | 2.23 |
| Residual | 17.55 | 17.32 | 17.78 |

***Figure S3*. Full regression model of gamma power and peak-frequency changes. Related to Figure 3.** (A) Random intercepts linear mixed regression model describing per-trial stimulus-induced percent gamma power changes from the baseline. Gray text indicates non-significant effects. (B) Same as (A), but describing per-trial stimulus-induced gamma peak frequency. For both models, see Methods.

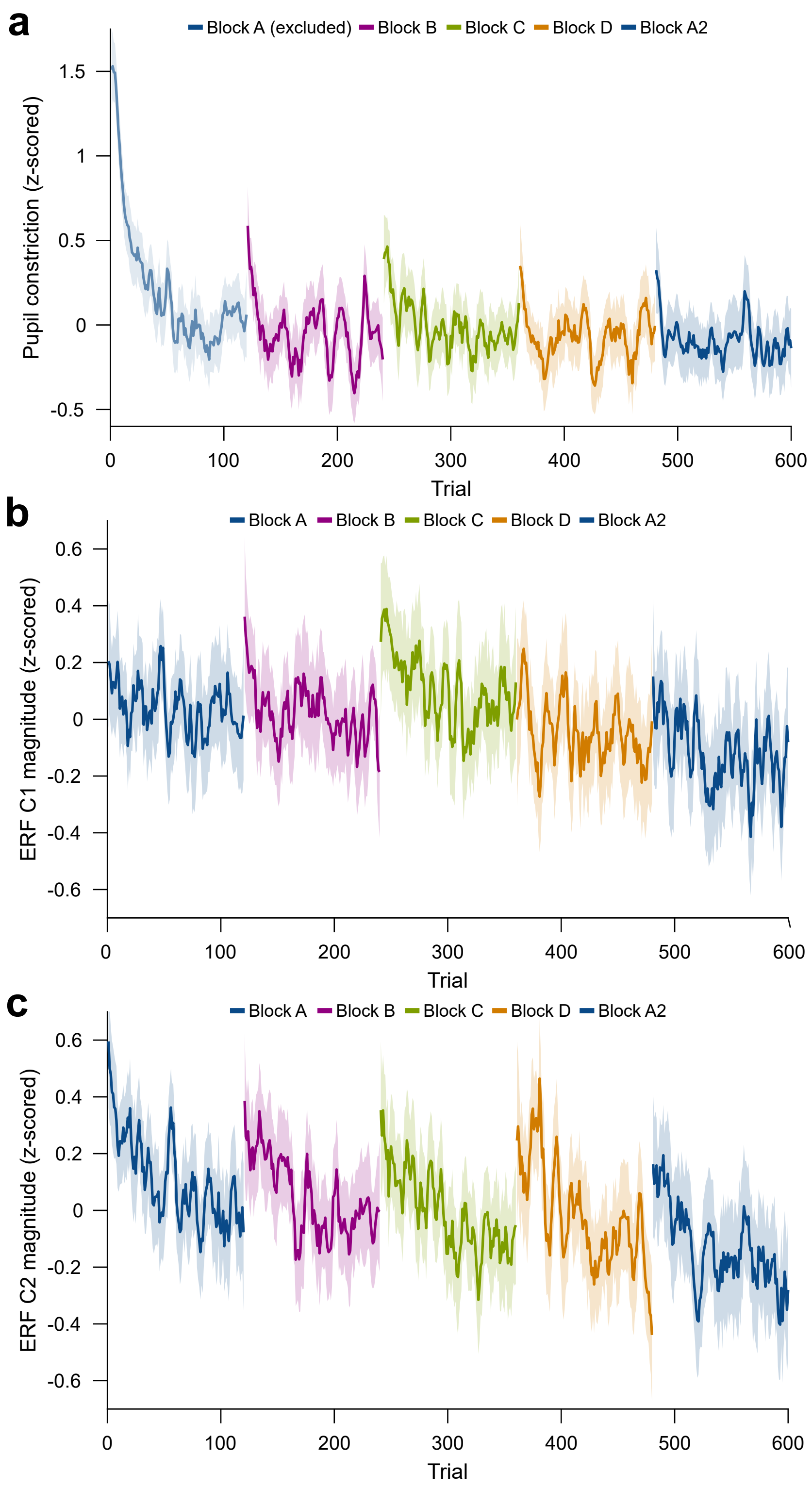


***Figure S4*. Per-trial changes in pupil constriction and ERF magnitude. Related to Figure 4.** (A) Per-trial average pupil constriction (the difference between pupil size before stimulus onset and 0.5 s-1.2 s after stimulus onset) as a function of trial number. The first block is confounded by ongoing pupil size adaptation to the changed level of ambient light and projector illumination at the beginning of the experiment and was excluded from pupil-size analyses. (B) Per-trial average magnitude of the ERF C1 component (55-70 ms post-stimulus). (C) Per-trial average magnitude of the ERF C2 component (90-180ms post-stimulus). For plots A, B, and C, values were z-scored within subjects. Average and 95% bootstrap confidence intervals were computed using a five-trial-wide running window within each block.

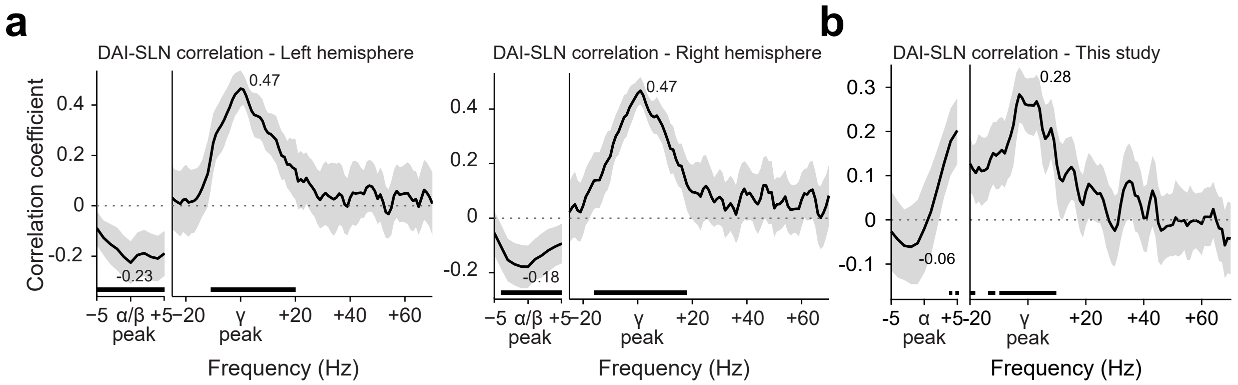


***Figure S5*. Conceptual replication of Michalareas et al.[18].** (A) Replotting of the data previously published in Fig. 6 of Michalareas et al.[18]. The graphs show the Spearman correlation coefficient, across pairs of brain areas, between a metric of GC asymmetry (DAI, per frequency) and a metric of the feedforward character of the corresponding anatomical projection (SLN). The DAI metric is the directed influence asymmetry index from brain area A to brain area B, defined as $DAI_{A-to-B} = [GC_{A-to-B} - GC_{B-to-A}]/[GC_{A-to-B} + GC_{B-to-A}]$). The SLN metric is the supragranular labeled neuron proportion, a graded anatomical metric (defined in the macaque) of the degree to which an anatomical projection is of feedforward character (i.e. originating in supragranular layers) [76]. DAI was positively related with SLN in the subject-defined gamma band, and negatively related to SLN in the subject-defined alpha/beta band. Lines show the average over subjects, and error bands represent standard error of the mean across subjects. Significance was computed using cluster-based nonparametric testing against zero [70]. (B) Similar analysis for the dataset recorded here, using the same seven brain areas. The same positive relation was found in the subject-specific gamma band. A similar negative relation is visible in the subject-specific alpha band, but did not reach significance. Lines show the average over subjects, and error bands represent the bootstrapped 95% confidence interval, significance was computed using a $t_{max}$-corrected permutation test. SLN values from Chaudhuri et al.[77].

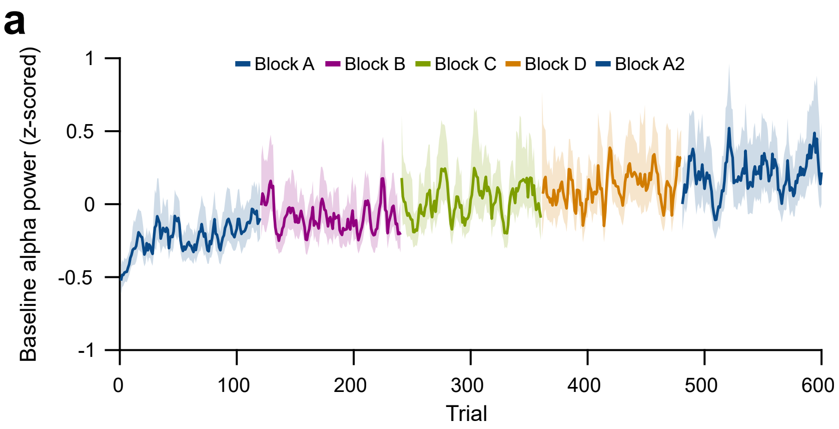


***Figure S6*. Per-trial changes in baseline alpha power.** (A) Per-trial average alpha power during the pre-stimulus baseline interval, z-scored within subjects. Average and 95% bootstrap confidence intervals were computed using a five-trial-wide running window within each block.
